# An increasingly efficient narrowband object-recognition channel along the ventral stream

**DOI:** 10.64898/2026.02.10.705069

**Authors:** Ajay Subramanian, Ekin Tünçok, Jan W. Kurzawski, Najib J. Majaj, Denis G. Pelli, Jonathan Winawer

## Abstract

How does the visual system recognize objects in natural environments? Here, we investigate the neural computations underlying robust object recognition along the ventral stream. We adapted psychophysical critical-band masking to fMRI, measuring BOLD responses along the ventral stream to natural images perturbed by bandpass noise. We found that, along this stream, the overall BOLD response to noise alone is sensitive to an increasingly wide range of stimulus spatial frequency, from 2 octaves in V1 to 5 octaves in ventral temporal cortex (VTC). However, when we assess the effect of the same noise on the accuracy of decoding scene images from the BOLD response, we obtain a different result: Recognition bandwidth is conserved along the ventral stream at about 2 octaves, close to the 1.5-octave behavioral channel. Though the recognition band is conserved, its noise tolerance increases steadily along the ventral stream, approaching behavioral levels in VTC. These findings suggest that V1 sets the bandwidth of the object-recognition channel, while downstream areas progressively denoise the signal, establishing the channel’s noise tolerance. This biological architecture—early channel isolation followed by successive denoising—may explain why human vision remains more robust than current machine vision.

## INTRODUCTION

People recognize gratings, letters, and natural objects using a 1.5-octave-wide spatial-frequency band (Campbell and Robson, 1968, Majaj et al., 2002, Solomon and Pelli, 1994, Subramanian et al., 2023).^1^ Noise in this band strongly impairs recognition, whereas noise outside it has little effect (Figure 1). The narrowness of this channel reflects a constraint of the visual system rather than a property of natural images (Figure S1) (Solomon and Pelli, 1994).

**Figure 1:**
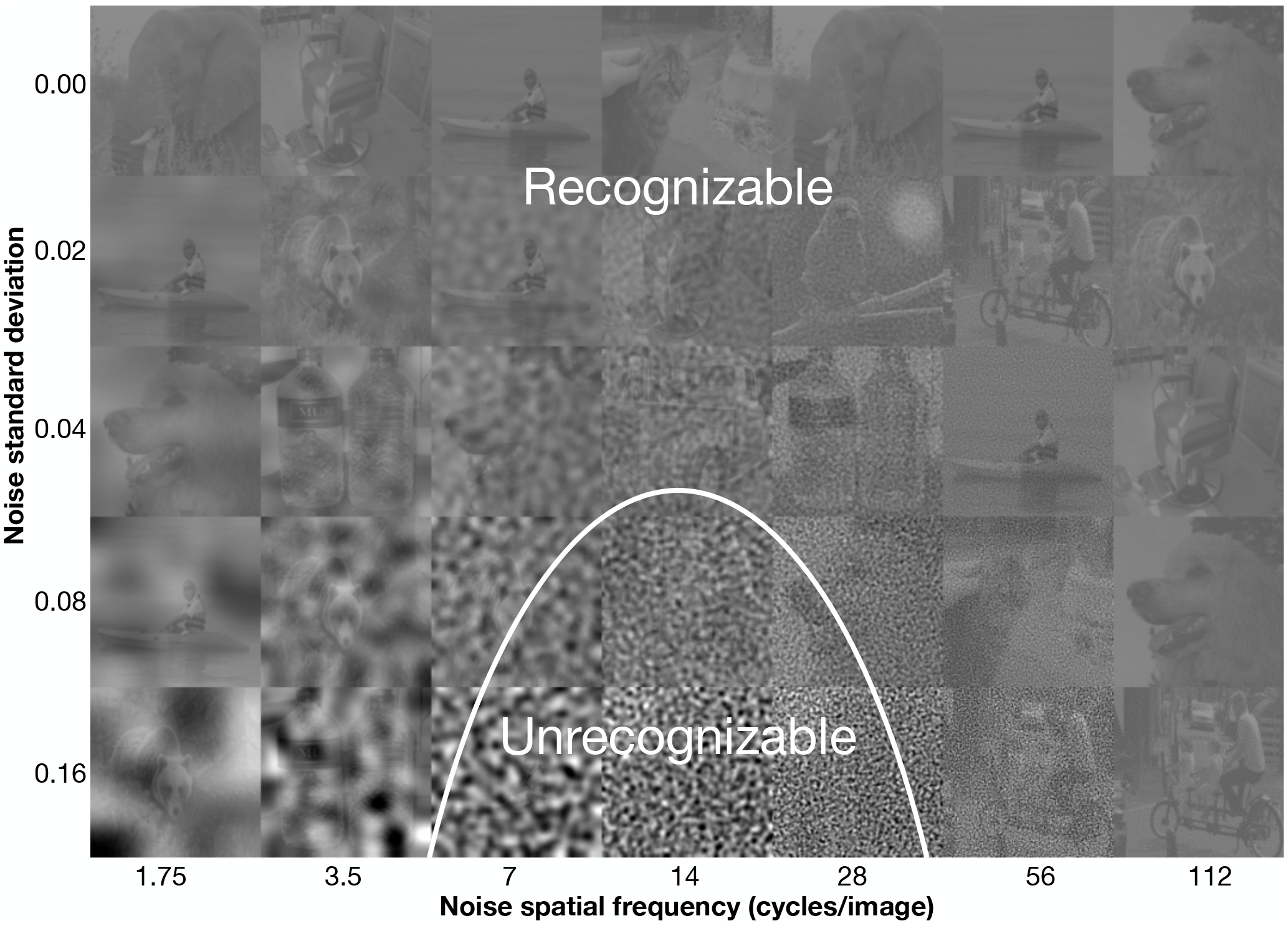
Critical-band masking and the 1.5 octave psychophysical object recognition channel. A demo of how critical-band masking reveals a spatial-frequency channel for object recognition. Each cell in the grid contains a sample grayscale, contrast-reduced (20%) image from ImageNet, perturbed with filtered Gaussian noise. Rows correspond to increasing noise standard deviation *σ* from top to bottom. Columns correspond to noise filtered within one-octave spatial-frequency bands centered at increasing frequencies from left to right. To visualize the channel, determine how far down each column you can go before being unable to recognize the object. These thresholds trace out an inverted-U-shape like curve shown in white, typically centered at 14 cycles/image, identifying the spatial frequencies most critical for recognition. The plot displays the noise threshold SD predicted by our spatial-frequency channel model as a function of spatial frequency, mirroring the perceptual sensitivity. The demo assumes a viewing distance of two times the width of the composite image.

By contrast, deep neural networks typically rely on a much broader spatial-frequency range—around four octaves (Subramanian et al., 2023). This broader tuning is associated with reduced robustness (Subramanian et al., 2023), including weaker shape bias (Geirhos et al., 2018a) and greater susceptibility to adversarial attack (Szegedy et al., 2013). Here, we investigate the physiological basis, in humans, of the narrow channel for object recognition. We adapt the psychophysical method of critical-band masking (Fletcher, 1940) to study the effect of stimulus noise on neural representations along the human recognition pathway: V1 to V2 to V3 to V4 to ventral temporal cortex (VTC).

We measure BOLD responses to bandpass noise images and to natural images perturbed by that noise (Figure S2). Across conditions, noise was filtered into seven one-octave bands centered at 1.75, 3.5, 7, 14, 28, 56, and 112 cycles/image and presented at five noise standard deviations: 0, 0.02, 0.04, 0.08, and 0.16 (in normalized intensity units).

We track the evolution of three measures along the ventral pathway and compare them with behavioral object recognition:

1. The *noise-response band* is the range of noise-stimulus frequencies in noise-alone images that elicit a criterion BOLD response.
2. The *recognition band* is the range of noise-stimulus frequencies in noise-plus-scene images that disrupt image decoding from the BOLD response.
3. *Noise tolerance* is the lowest noise power, across all frequency bands, that prevents image decoding.

The *noise-response band* specifies the frequencies that drive neural activity, the *recognition band* specifies the frequencies that interfere with scene decoding, and *noise tolerance* quantifies how efficiently the system (classification of BOLD response to stimulus) decodes scenes in noise.

## RESULTS

Using critical-band masking, we measured three properties of neural representations along the ventral stream: the noise-response band, the recognition band, and noise tolerance.

### 1. Along the ventral stream, the BOLD response is sensitive to a progressively broader range of noise frequencies

We first measured the response of each visual area to bandpass noise alone. The amplitude of the BOLD response depends strongly on the spatial frequency of the noise. We fit a spatial-frequency channel model to these responses separately for each visual area.

From these responses we define the *noise-response band* as the range of stimulus frequencies for which the BOLD response exceeds a criterion level. Figure 2A shows the noise-response bands for each visual area. Figure S4 shows the bands for all observers. The noise-response band broadens systematically along the ventral stream. In V1 the band is narrow, about 2 octaves wide. In successive visual areas the band becomes progressively wider, reaching about 5 octaves in VTC. This is consistent with Ha et al. (2025), who also found increasing bandwidth from V1 to V3 (but see Aghajari et al. (2020)).

**Figure 2:**
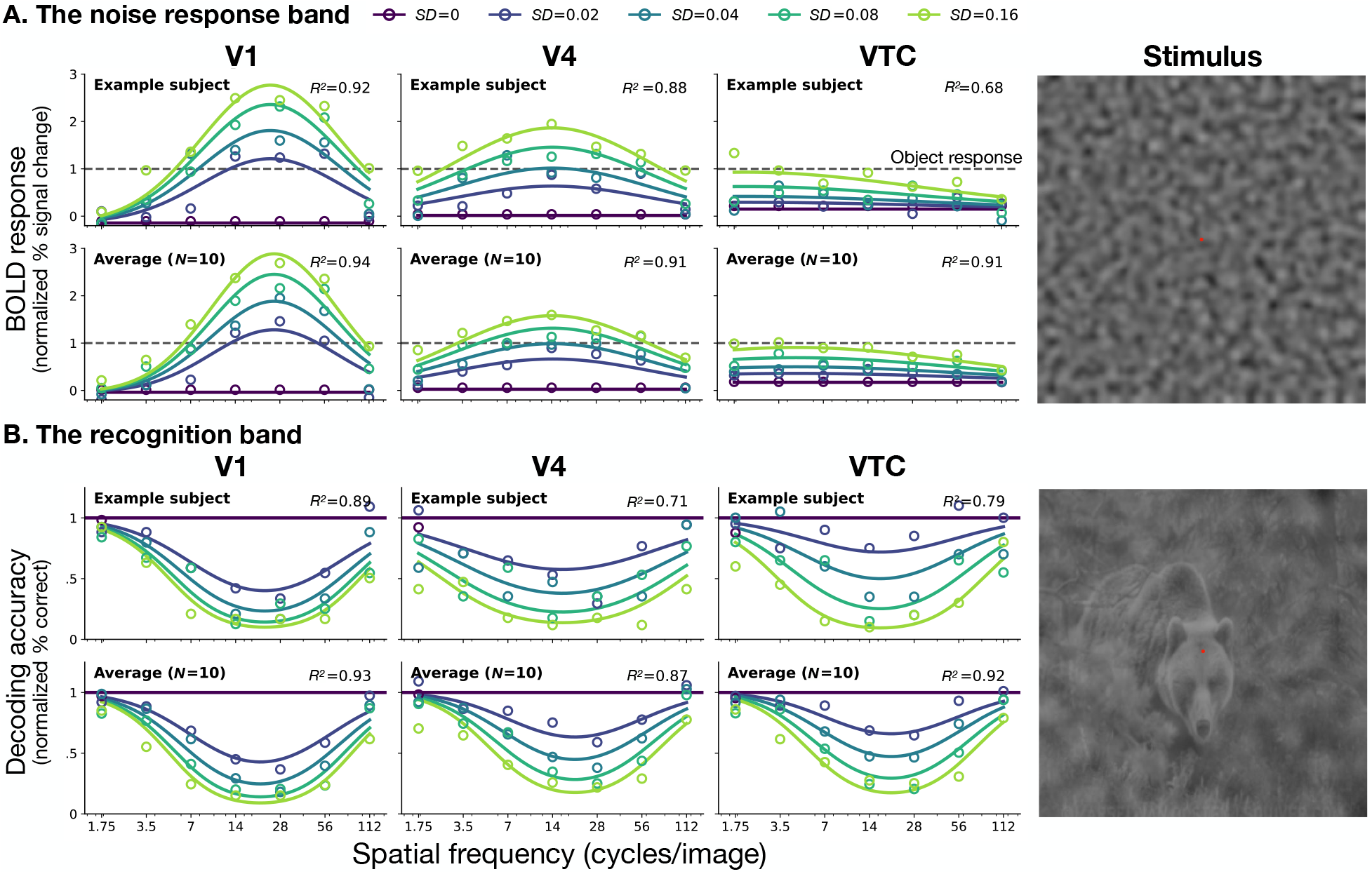
Divergent tuning of noise-response and object recognition bands along the ventral stream. **A. The noise-response band**. BOLD response amplitude (% BOLD signal change) to bandpass noise alone, normalized by scene-alone response, as a function of spatial frequency. Rows show data and fits for an example observer (top) and the average across observers (bottom) in areas V1, V4, and VTC. Color indicates noise standard deviation *σ*. Dashed horizontal lines represent the response amplitude to the scene alone in each area (=1, normalized). **B. The recognition band**. Accuracy of object decoding from noisy images, normalized by decoding accuracy for zero-noise images, as a function of noise spatial frequency. Rows correspond to the individual (top) and average (bottom) observer for regions V1, V4, and VTC. Solid horizontal lines indicate baseline decoding accuracy for noiseless images (*σ* = 0). Circles represent measured data points; solid curves represent predictions from the fitted spatial-frequency channel models, with *R*^2^ values indicating goodness-of-fit for each ROI. The images on the right shows a sample stimulus from the experiment used to measure each band.

The band also shifts toward lower spatial frequencies. Thus, as signals progress along the ventral stream, neural responses become sensitive to a broader range of spatial frequencies, with increasing weight on lower frequencies.

Comparing these responses to the response amplitude to natural images (dashed lines), we see a shift from preferring noise in V1 to preferring images of natural scenes in VTC.

### 2. Although sensitivity to noise broadens along the ventral stream, the recognition bandwidth is conserved

To determine whether the broadening of the response to noise implies a broadening of the effect of the noise on recognition, we applied an image classifier to the BOLD response to noisy scenes. This yielded a *recognition band* for each visual area (Figure 2B). Figure S5 shows recognition bands for all observers in all visual areas. The patterns in both the noise-response and recognition bands are also conserved across eccentricity (Figure S7, S8).

In contrast to the widening noise-response band, the recognition band is conserved along the ventral stream. For example, in V1 the effect of noise on decoding is approximately mirror-symmetric to the response to noise alone, whereas in VTC the two patterns are strikingly (Hung et al., 2005, Rust and DiCarlo, 2010).

We summarize the divergence by calculating the bandwidth of the noise-response band and the recognition band for each area (Figure 3, left). The noise-response bandwidth increases monotonically from 2 octaves in V1 to 5 octaves in VTC. In contrast, the recognition bandwidth is conserved at 2 octaves, close to the 1.5 octave band measured psychophysically.

**Figure 3:**
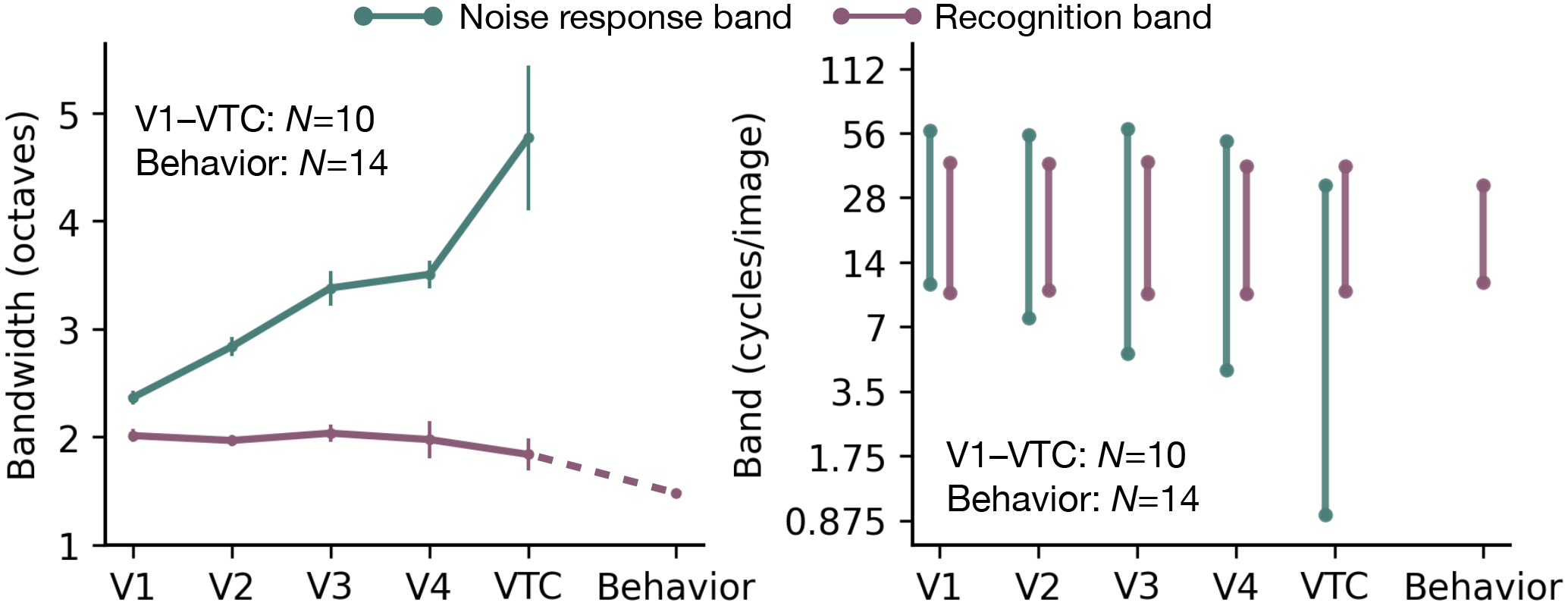
Divergence between noise-response and recognition bands. **Left:** Channel bandwidth is plotted for each visual area, V1 to VTC, and behavior. **Green line** shows the bandwidth of the noise-response band (defined as the full-width at half max, FWHM, of the fitted Gaussian tuning), which widens progressively from 2 octaves in V1 to 5 octaves in VTC. **Purple line** shows the bandwidth of the recognition band (defined as the width at twice the minimum power threshold), which remains conserved at 2 octaves in all areas, close to the behavioral 1.5 octave recognition bandwidth. Data points represent the mean across subjects (*N* = 10 for fMRI, *N* = 14 for psychophysics), and error bars indicate ±1 SEM. For behavior, the error bar is smaller than the dot size. **Right:** Frequency range of each band for each visual area. From V1 to VTC, the *noise-response* band grows broader and shifts lower. *Recognition* bandwidth and center frequency are conserved from V1 to VTC at values similar to the behavioral recognition band. Note that the ventral-stream bands in this subplot are from model fits to the averaged data across subjects i.e., the bands shown in bottom panels of Figure 2A,B.

The two bands also have different positions. The noise-response band becomes wider and shifts toward lower spatial frequencies from V1 to VTC. The recognition band, however, maintains both its bandwidth and its preferred frequency (19 cycles/image) across all areas, closely matching the behavioral recognition band (Figure 3, right).

### 3. Along the ventral stream, noise response weakens and scene recognition becomes more noisetolerant

Next we quantify how sensitivity to noise changes along the ventral stream. We define the *noise threshold* as the noise power required for noise alone to elicit a BOLD response equal to that of the scene alone. We define *noise tolerance* as the noise power required to halve the accuracy of decoding scene images.

Both measures increase along the ventral stream (Figure 4). The noise threshold (green line) rises steeply from V1 to VTC—a 2^8^-fold increase—indicating that downstream areas require much stronger noise to produce a response comparable to that of scenes. This is presumably related to the fact that higher visual areas are sensitive to higher-order structure that is absent in the noise stimuli (Freeman et al., 2013, Lieber et al., 2024, Ziemba et al., 2016). Successive areas respond less to noise relative to scenes, i.e. they *denoise* the input. Further evidence of denoising comes from a superposition test: the response to noise is greatly reduced by the presence of scenes in VTC but not in V1 (Figure S3). In the same figure, the reduction is especially large in the spatial frequencies within the recognition band.

**Figure 4:**
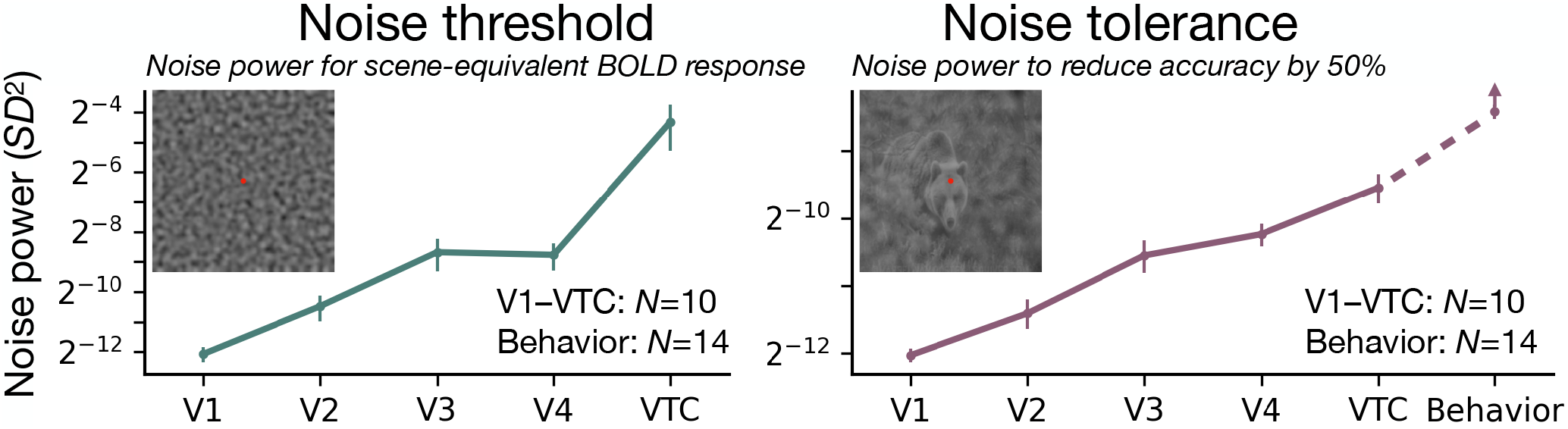
The noise threshold and tolerance both increase along the ventral stream. **Left:** Noise power *σ*^2^ that elicits a BOLD response equal to that of the scene alone. For each area, the noise threshold is calculated for the most effective frequency band. Later areas require more noise to reach scene-equivalent response, with VTC’s threshold being 2_8_ × higher than V1’s, indicating a much improved signal-to-noise ratio (SNR). For a complementary analysis in terms of response amplitude instead of threshold, see S6. **Right:** Noise power required to halve (50%) decoding accuracy in each visual area. Noise tolerance increases by 2^3^ × from V1 to VTC, approaching the tolerance measured in behavioral object recognition, indicating that higher-level object representations tolerate noise levels that disrupt early visual areas. Upward arrow on the behavioral error bar accounts for the possibility that the behavioral sensitivity would be even greater if the behavioral task had used the same number of categories (10 vs 16) and exemplars per category (1 vs 30) as the fMRI task. Data points show the mean across subjects (fMRI: *N* = 10; psychophysics: *N* = 14). Error bars indicate ±1 SEM.

The noise tolerance (purple line) also increases but less steeply, 2^3^-fold from V1 to VTC, indicating that the high-level representations of scenes are more robust to noise. In VTC, the noise tolerance of image decoding approaches that of behavioral recognition (dashed purple line).

## DISCUSSION

Our results reveal how the visual system manages signal and noise along the ventral stream. Although the BOLD signal responds to an ever broader band of noise frequencies, the critical band that disrupts recognition remains narrow (near behavior), while noise tolerance increases towards behavior. Thus, as signals progress from V1 to VTC, neural responses become less sensitive to noise, enabling increasingly noise-tolerant image decoding. Together, these results suggest that object recognition improves along the ventral stream not by narrowing the object channel but by increasing tolerance to noise.

### Estimating a linear filter in a nonlinear model of the brain

Recognition and the brain are nonlinear, but the ventral stream begins with V1, whose responses are less nonlinear, as shown by a superposition test (Figure S3). Unlike V1, where noise and scene responses are close to additive, in VTC, the noise response is strongly suppressed by the scene.

Psychophysically, analysis of critical-band masking typically assumes a linear filter followed by a nonlinear recognition process. This assumption has been validated in object recognition experiments by superimposing two noises with non-overlapping spectra, finding a near additive effect on the energy threshold (Solomon and Pelli, 1994).

### Robustness to noise in humans and machines

Using only 2 octaves for recognition is sub-optimal in an information-theoretic sense. An ideal observer would achieve the lowest possible signal energy threshold by utilizing all available spatial frequencies (Geisler, 2011, Neri, 2015, Solomon and Pelli, 1994). For many tasks, deep neural networks, like ideal observers, use a broad range of frequencies, but this broad tuning makes them vulnerable to attack (Subramanian et al., 2023). By strictly limiting the readout to a narrow band, the human visual system seems to sacrifice information to achieve greater robustness.

Most CNNs lack the narrow recognition channel of human vision (Subramanian et al., 2023). However, recent foundation models (large models trained with large amounts of data) perhaps perform differently. These networks, e.g., aligned with language (Radford et al., 2021) or adversarially-trained (Madry et al., 2017), show improved robustness to noise. The improved robustness might make them more similar to human behavior, but they achieve this through a mechanism distinct from the human brain (Chen et al., 2020, Madry et al., 2017). Their training objectives — whether through contrastive learning or adversarial defense — explicitly force the model to map a noisy image to the same point in feature space as its noise-free counterpart. This pressure drives the network to suppress noise entirely, effectively making the system blind to the perturbation. In the brain, the response to noise is still present in VTC, although much weaker than in V1. To build truly-brain-like models, our aim should be to build systems that tolerate noise instead of being blind to it. The brain does not completely suppress the “fog” to see the “car”; it represents both.

### Effects of Estimating a linear filter in a nonlinear model of the brain120 size

Our experiment was conducted at a fixed stimulus size. Our expectation is that if the stimulus size changed, the critical bandwidth in octaves would be unchanged, as this has been found for critical-band masking of letters (Majaj et al., 2002). Effects of stimulus size on the most effective masking frequency are more complicated and have been shown to follow a 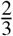 power law, being neither constant in cycles/deg nor in cycles/image. Recent work provides preliminary evidence that these observations extend to natural images (Yancy et al., 2026, VSS, *unpublished*).

### Effects of task and attention

In our study, observers performed a fixation task unrelated to the scene images. The recognition objective was only imposed during analysis, via image decoding. Under an active recognition task, there might be top-down modulation of ventral-stream representations (Harel et al., 2014, Maunsell and Treue, 2006), leading to more accurate decoding (Kay et al., 2015). However, the improvement of noise tolerance from V1 to VTC is independent of attention (Pratte et al., 2013). Moreover, since the recognition bandwidth along the ventral stream is already near behavior, we can conclude that attention is not needed to achieve a narrow recognition channel.

## CONCLUSION

Perhaps the most surprising result is the divergence between how the brain responds to noise alone and how noise affects the response to scenes. The noise-response bandwidth grows from 2 to 5 octaves along the ventral stream, becoming less and less like the behavioral bandwidth. In contrast, the recognition bandwidth is conserved along the ventral stream and is similar to the 1.5-octave behavioral bandwidth. The sensitivity to noise alone decreases greatly along the ventral stream, 2^8^-fold lower in VTC than in V1. The tolerance of image decoding to stimulus noise improves by 2^3^ along the ventral stream. Thus, the ventral stream achieves highly robust object recognition not by filtering out the noise entirely, but by decreasing noise sensitivity, increasing noise tolerance, and maintaining a narrow channel for object recognition.

## RESOURCE AVAILABILITY

### Lead contact

Requests for further information and resources should be directed to Ajay Subramanian.

### Materials availability

This study did not generate new materials.

### Data and code availability

All data and code will be released upon publication. Reviewers can access stimuli, code, data at https://osf.io/mxatg/overview?view_only=11cc8e5458c34866b91eba9f58d7cb31

## ACKNOWLEDGMENTS

Thanks to Alec Marantz for suggesting we extend our earlier psychophysical critical-band masking study to fMRI. Thanks to Eero Simoncelli for suggesting we look at the progression of noise tolerance along the ventral stream. Thanks to Eero Simoncelli, Michael Landy, Grace Lindsay, and Laura Suciu for their comments and suggestions on an earlier draft. Thanks to Furkan Özçelik for early discussions. This research was supported by the National Institutes of Health (R01 EY027401 and R01 EY033628 to JW; R01 EY031446 and RF1 NS127122 to NJM; P30 EY013079 Core Vision Grant), pilot grants from the New York University Center for Brain Imaging, ASPIRE Precision Medicine Research Institute Abu Dhabi VRI-20-10, and software from University of Minnesota Center for Magnetic Resonance Research. This work was supported in part through the NYU IT High Performance Computing resources, services, and staff expertise.

## AUTHOR CONTRIBUTIONS

AS, NJM, DGP, and JW conceptualized the project. AS, ET, NJM, DGP, and JW acquired funding. ET, JWK, AS, and JW designed the experiment. ET collected, pre-processed and prepared fMRI data for modeling analysis. AS and JW performed modeling analysis. AS, DGP, JW, NJM, and ET wrote and edited the paper.

## DECLARATION OF GENERATIVE AI AND AI-ASSISTED TECHNOLOGIES

During the preparation of this work, the authors used AI tools, including ChatGPT, Google Gemini, Cursor and Google Antigravity, for code generation and paper writing. After using these tools and services, the authors reviewed and edited the content as needed, and we take full responsibility for the content of the publication.

## SUPPLEMENTAL INFORMATION INDEX

Figures S1-S8, their legends, and captions.

## METHODS

## 1 STUDY PARTICIPANT DETAILS

Data were collected from 10 healthy human volunteers (5 female, 5 male). All participants had normal or corrected-to-normal visual acuity. Participants provided written informed consent in accordance with the experimental protocol approved by the New York University Institutional Review Board. Age, sex, and gender were not considered as variables in the study design.

## 2 METHOD DETAILS

### 2.1 Apparatus

Experimental stimuli were generated on an Apple iMac MATLAB (R2023) and Psychtoolbox-3 (Kleiner et al., 2007) and displayed on a ProPixx DLP LED projecter (VPixx Technologies Inc., Saint-Bruno-de-Montarville, QC, Canada). The display luminance was linearized (gamma-corrected) so that pixel intensity values were proportional to emitted luminance. This ensured that the additive Gaussian noise we generated in image intensity space was not distorted by display nonlinearities that could otherwise alter its distribution. Participants viewed the projected screen (60 x 36.2 cm; 1920 x 1080 resolution; refresh rate of 60 Hz) through an angled mirror mounted on the scanner’s head coil. The viewing distance from the eye to the mirror to the screen was 86.5 cm. Participant responses were collected using a 4-button fiber optic response box (Current Designs). Participants could use any of the four buttons to respond. Head pads were used inside the coil to stabilize participants head and reduce head motion and discomfort.

### 2.2 Stimuli

Stimuli for the main experiment were generated using the same image set and preprocessing pipeline as Subramanian et al. (2023). Briefly, we started from grayscale versions of natural images drawn from the 16-class ImageNet subset (Deng et al., 2009, Geirhos et al., 2018b) and selected 10 exemplar images from 10 distinct superordinate categories (bird, dog, cat, bear, elephant, truck, boat, chair, bottle, bicycle). Images were first resized to 256 × 256 pixels, center-cropped to 224 × 224 pixels (the standard ImageNet preprocessing protocol), and then upsampled to 512×512 pixels for presentation. All images were converted to grayscale and normalized to have low contrast by scaling their pixel intensities to 20% of their original root-mean-square (RMS) contrast about a midgray level (intensity 0.449 on a [0,1] scale). This ensured that adding high-strength noise would not produce substantial pixel clipping at floor or ceiling.

Bandpass noise was added to these low-contrast images following the critical-band masking procedure described in Subramanian et al. (2023). On each trial, we sampled a Gaussian white-noise field and, for non-zero noise conditions, filtered it into one of seven octave-wide spatial-frequency bands using a Laplacian pyramid decomposition (Burt and Adelson, 1987). The seven bands corresponded to successive levels of the pyramid and were approximately centered at 1.75, 3.5, 7, 14, 28, 56, and 112 cycles per image, each spanning one octave (a factor of two in spatial frequency). For each band, the filtered noise was rescaled so that its RMS contrast matched a desired noise strength and then added to the low-contrast scene. We used five noise strengths (noise standard deviations of 0, 0.02, 0.04, 0.08, and 0.16, in units of normalized image intensity). The zero-noise condition (SD = 0) yielded low-contrast images without added noise. After addition of noise, pixel values were clipped to the [0,1] range if necessary.

In addition to noise-masked scenes, we also generated noise-only stimuli by adding bandpass noise (constructed as above) to a uniform midgray image (intensity 0.449). This produced a matched set of bandpass noise patterns at each spatial frequency and noise strength, without any overlaid scenes. All stimuli were presented centrally on a uniform gray background, and the same set of object and noise conditions was used across all fMRI runs.

### 2.3 Experimental Protocols

#### Main experiment

Participants completed 11, 13 or 15 scans of the main event-related fMRI experiment (Table 1). Each image, created as described above, was presented for 3 sec with 1 sec off time between two consecutive image presentations. We adapted this protocol from previous work that used event-related presentation of natural images and elicited reliable responses from visual cortex (Allen et al., 2022). Participants viewed 87 images on each scan session, totaling up to the following numbers of total image presentation across the experiment:

**Table 1.** Trial counts per participant. Each cell shows unique stimuli × repetitions.

| SID | Runs | Total trials | Scenes in noise | Noise only | Scene only | Blank |
| --- | --- | --- | --- | --- | --- | --- |
| 145 | 13 | 1133 | $280 \times 3$ | $28 \times 3$ | $10 \times 19$ | $1 \times 19$ |
| 135, 136 | 11 | 1307 | $280 \times 3$ | $28 \times 9$ | $10 \times 19$ | $1 \times 25$ |
| 114, 127, 141, 146, 147, 151, 185 | 15 | 1305 | $280 \times 3$ | $28 \times 9$ | $10 \times 19$ | $1 \times 23$ |

Throughout the experiment, participants were asked to fixate at the center of the display. They performed a color change detection task at the fixation dot by making a button press. Each scan lasted 6 min 36 seconds (396 TRs).

#### Retinotopic mapping

Participants completed six scans of a retinotopic mapping experiment adapted from Allen et al. (2022). We used two different types of stimulus aperture, moving bars (three scans) and moving wedges and rings (three scans). We placed colorful texture patterns inside the apertures composed of different types of visual objects. These images were randomly drawn from the image dataset previously used for retinotopic mapping (Allen et al., 2022, Benson et al., 2018, Kriegeskorte et al., 2008). Mapping stimulus flickered at 3Hz on a given location in visual field. The apertures covered up to 12.4° in eccentricity. Participants were asked to fixate throughout the experiment as the retinotopic mapping stimulus moved across the visual field. They performed a color change detection task at fixation, where they were asked to make a button press when the fixation cross changed color. Each scan lasted 5 mins (300 TRs). Runs with moving bars and moving wedges and rings were interleaved.

#### Functional localizers for mapping ventral temporal cortex

Participants completed four scans of a functional localizer experiment targeting different category-selective visual cortical areas. We used an experimental protocol based on previous work that successfully delineated category-selective areas of ventral temporal cortex (https://github.com/VPNL/fLoc) (Stigliani et al., 2015). Images from a total of five different categories, bodies, characters, faces, objects, places, were shown to the participants in an event-related fMRI protocol. Body images consisted of bodies with removed heads and limbs, character images consisted of pseudowords, faces were those of adults, object images consisted of different cars, and place images were different houses. These images were overlaid on a phase-scrambled gray background that extended 10.5° in eccentricity. Each scan contained 30 distinct image presentations with each image presented for 500 ms, interspersed with blank periods. Images from different categories were shown in a randomized order in each scan. Participants were asked to fixate throughout the experiment. Each scan lasted 3 min 48 seconds (240 TRs).

#### MRI data acquisition

We collected functional and anatomical MRI data using a 3T Siemens Prisma scanner with a 64-channel head/neck coil at NYU’s Center for Brain Imaging. All functional time series data (from the main experiment, retinotopic mapping experiment, functional localizer experiment) were registered to the same anatomical scan collected during the retinotopic mapping experiment. T1-weighted anatomical images were acquired with the following scan parameters: MPRAGE: TR = 2400 ms, TE = 2.2 ms, 0.8 mm isotropic voxels, flip angle = 8°. For the main experiment, we collected 11, 13 or 15 runs (Table 1) of functional echo-planar images (EPIs) from each participant across two separate sessions. Prior to the acquisition of functional echo-planar images (EPIS), we first acquired distortion maps with opposite phase-encoding directions (AP and PA) to correct for susceptibility-induced distortions in the functional images. Functional runs in each session were acquired using a CMRR multiband EPI sequence (TR = 1000 ms, TE = 37.6 ms, 2 mm isotropic voxels, flip angle = 68°, multiband acceleration factor = 6 (Feinberg et al., 2010, Moeller et al., 2010, Xu et al., 2013)). Functional EPIs were acquired with the same scan parameters across all three experiments. Three experiments differed in the number of TRs (slice dimensions 104*x*104*x*66; the main experiment: 396 TRs, retinotopic mapping experiment: 300 TRs, functional localizer experiment: 240 TRs).

### 2.4 Data preparation

#### Preprocessing

DICOM files were anonymized and defaced with pydeface before being transferred from the scanner to the data server. The original DICOM files were next converted to NIfTIs and organized according to Brain Imaging Data Structure (BIDS) (Gorgolewski et al., 2016) conventions using heudiconv. For preprocessing, we used fMRIPrep 20.2.7 (Esteban et al., 2019). First, T1-weighted anatomical images were bias field corrected and skull stripped. followed by tissue segmentation (CSF, white matter, gray matter) using FAST with both T1w and T2w inputs. Cortical surface reconstruction was performed with FreeSurfer’s *recon-all* (Dale et al., 1999). For functional data, we generated a reference volume, applied skull stripping, and corrected for susceptibility distortions using the AP and PA field maps. The corrected functional reference was coregistered to the anatomical image using *bbregister* (Greve and Fischl, 2009). We estimated head motion parameters relative to this reference and applied slice-timing correction with AFNI’s *3dTshift* (Cox and Hyde, 1997). The slice-time corrected data were then resampled to anatomical space in a single interpolation step that combined all spatial transformations (motion correction, distortion correction, and coregistration). Finally, the preprocessed BOLD data were projected onto each participant’s native cortical surface *(fsnative)*, and all subsequent analyses were conducted in this surface-based space.

#### Population receptive field (pRF) model

We ran a circular 2D Gaussian population receptive field (pRF) model on each vertex timeseries data from the retinotopic mapping experiment in each participant’s native brain surface. We used the *vistasoft* software (github.com/vistalab/vistasoft, Vista Lab, Stanford University), with a custom wrapper function (github.com/WinawerLab/prfVista, New York University to deploy the pRF models for each participant. Each vertex pRF was parameterized by a center (*x, y* in deg) and a size (*σ*, one standard deviation of the Gaussian, in deg) (Dumoulin and Wandell, 2008). The responses of each vertex to the moving retinotopic mapping stimuli were predicted by multiplying the Gaussian pRFs with the binarized stimulus aperture in time, then convolving it with a hemodynamic response function to account for the delay in BOLD response. For each vertex, pRF models were fit to the averaged timeseries data from three scans of two separate retinotopic mapping stimuli (moving bars and moving wedges-rings). For each vertex, the model estimated three parameters (*x, y, σ*) by minimizing the sum of squared errors between the measured responses from the averaged timeseries data and the predicted responses from the model. The fitting procedure used a coarse-to-fine approach with HRF optimization: an initial grid search over candidate pRF parameters, followed by iterative minimization with temporal decimation, then without decimation, and finally a search to optimize the hemodynamic response function parameters jointly with the pRF parameters. Goodness-of-fit was assessed using the coefficient of determination, 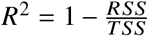, reflecting the proportion of variance in the measured timeseries explained by the pRF model.

#### Defining V1, V2, V3, and V4

We used Neuropythy v0.12.11 (Benson and Winawer, 2018) to delineate the early visual cortical ROI boundaries on the native brain surface of each participant. First, we visualized the eccentricity and polar angle solutions of each vertex pRF on the cortical surface, centered at the occipital pole. Then following the criteria laid out in previous papers for V1-V3 (Benson et al., 2022) and V4 (Kurzawski et al., 2025, Winawer and Witthoft, 2015) based on reversals of polar angle and eccentricity solutions, we manually drew the boundaries of these areas.

#### General linear model for functional localizers

To define ventral temporal cortex in each participant, we estimated category-selective responses in visual cortex using the functional localizer experiment defined above. We ran GLMsingle (Prince et al., 2022) on the preprocessed time series data from the four localizer runs. A design matrix was constructed with five predictors corresponding to each stimulus category (faces, bodies, cars, houses, words). Beta weights were estimated for each category at each vertex and converted to percent signal change.

#### Defining ventral temporal cortex

We used FreeSurfer v7.3.2 (Fischl, 2012) to manually define a ventral temporal cortical (VTC) area in each participant’s native inflated brain surface. To define VTC, we used a combination of anatomical and functional criteria laid out in previous work (Kay and Yeatman, 2017, Rosenke et al., 2020). First, for each participant, we computed category-selective contrast maps by subtracting the mean response (percent signal change) to four categories from the response to the remaining category (e.g., faces vs. bodies, cars, houses, and words). This yielded five contrast maps per participant, one for each category. To guide ROI definition, we created an aggregated contrast map combining three category contrasts: faces vs. others, bodies vs. others, and houses vs. others. For each contrast, vertices exceeding a threshold of 1% were labeled, and the three thresholded maps were summed to create a single overlay highlighting vertices with category-selective responses. These aggregated contrast maps were used in combination with anatomical landmarks that coincide with face-, body-, and place-selective ventral cortical areas: the midfusiform sulcus (MFS), which separates face-selective cortex on its lateral bank from word-selective cortex on its medial bank (Grill-Spector and Weiner, 2014, Weiner et al., 2014); the occipito-temporal sulcus (OTS), which marks the lateral extent of body-selective cortex (Peelen and Downing, 2005, Schwarzlose et al., 2005); and the collateral sulcus (CoS), which marks the medial extent of place-selective cortex (Epstein and Kanwisher, 1998, Weiner et al., 2018). VTC was defined in each hemisphere as the cortical region bounded laterally by the OTS, medially by the CoS, posteriorly by the posterior transverse collateral sulcus, and anteriorly by the anterior tip of the mid-fusiform sulcus.

#### General linear model

To model the responses of each vertex on every trial in our main experiment, we ran a general linear model on each vertex time series data using GLMsingle (Prince et al., 2022). GLMsingle integrates three techniques to improve the accuracy of single-trial beta estimates: (1) for each voxel, a custom hemodynamic response function is selected from a library of candidate functions, (2) cross-validation is used to derive noise regressors from voxels unrelated to the experimental paradigm, and (3) ridge regression is applied on a voxel-wise basis to regularize beta estimates for closely spaced trials.

The single-trial GLM was run on the preprocessed time series data of each participant concatenated across two fMRI sessions. There were 319 unique stimulus conditions: 11 image categories (10 objects + 1 blank) × 29 conditions (28 noise conditions + 1 no-noise condition). The number of trials varied across participants (see Table 1). A design matrix was constructed for each run with columns corresponding to each stimulus presented during that run. For each predictor column, a 1 was entered for each TR when the stimulus was on screen.

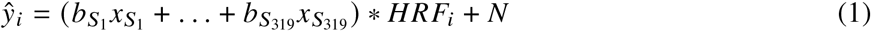

where *ŷ*_*i*_ represents the predicted time series of vertex *i*, the *b* terms are the coefficients (beta weights), and the *x* terms are indicator variables (either 0 or 1). The subscripts denote the stimulus condition, with *S*_1_ to *S*_319_ corresponding to the 319 unique stimulus conditions. All stimulus predictors were convolved with a voxel-specific hemodynamic response function (*HRF*_*i*_) selected from a library of candidate functions. *N* represents nuisance variables (polynomial regressors for detrending low-frequency fMRI drift and task-irrelevant noise regressors identified by GLMsingle). Estimated beta weights were converted to percent BOLD signal change by dividing by the mean signal intensity of each vertex. The resulting beta estimates were spatially smoothed on the cortical surface using FreeSurfer’s `mri_surf2surf` with a 3 mm full-width at half-maximum (FWHM) Gaussian kernel (Dale et al., 1999).

### 2.5 Analysis

#### Notation

To formalize our analysis, we introduce the following notation. Let *s* ∈ {1, …, *N*_*sub j*_} denote a subject and *r* ∈ {*V*1, *V*2, *V*3, *V*4, *VTC*} denote a region of interest (ROI). The stimuli are defined by object category *c* ∈ {1, …, *C*}, noise spatial frequency *f* ∈ *F*, and noise contrast (standard deviation) *σ* ∈ Σ.

For a given subject and ROI, let **y**_*t*_ ∈ ℝ^*v*^ represent the vector of GLM beta weights for trial *t*, where *V* is the number of voxels in the ROI. We denote the set of trials corresponding to a specific condition as *T* (*c, σ, f*). For noise-only trials, the category is undefined (or null), denoted as *T*_noise_(*σ, f*). For object trials, we denote the true category of trial *t* as *c*_*t*_.

#### Postprocessing

Our goal is to transform the trial-wise activation maps **y**_*t*_ into two summary matrices per subject and ROI: a *noise-response matrix* (quantifying the response band) and an *object-decoding accuracy matrix* (quantifying the recognition band).

##### 1. Voxel selection

Before generating matrices, we filtered voxels to ensure valid retinotopic coverage and responsiveness. For all ROIs, we included only voxels *v* with population receptive field eccentricities ecc_*v*_ ∈ [0°, 6°]. For VTC, we applied an additional inclusion criterion, selecting only voxels where the GLM variance explained 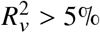.

##### 2. Noise-response matrix

To measure the response band, we computed the average magnitude of the BOLD response to noise alone. For each noise condition defined by frequency *f* and contrast *σ*, we averaged the activity across all voxels and all noise-only trials:

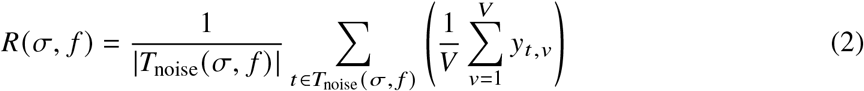

This yields a scalar response value for each condition, resulting in a matrix *R* ∈ ℝ^| Σ|×|*F*|^ for each ROI.

##### 3. Object-decoding accuracy matrix

To measure the recognition band, we evaluated how well object identity could be decoded from neural patterns under critical-band masking. We used a correlation-based nearest-centroid classifier.

First, we computed a *prototype µ*_*C*_ for each object category *c* by averaging the voxel patterns of all noiseless trials (*σ*= 0):

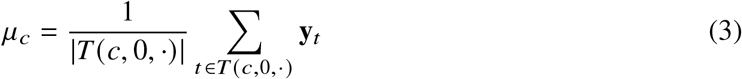

For decoding a test trial *t* (with noise parameters *σ, f*), we calculated the Pearson correlation ρ between the trial pattern **y**_*t*_ and each category prototype *µ*_*k*_. The predicted category *ĉ*_*t*_ was the one maximizing the correlation:

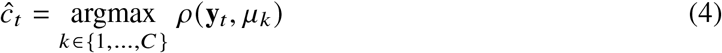

When decoding noiseless trials (where the test trial would otherwise be part of the prototype), we employed a leave-one-out procedure: the test trial was excluded from the calculation of its corresponding category prototype 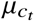 to prevent circularity.

Finally, we computed the decoding accuracy *A*(*σ, f*) as the proportion of correctly classified trials for each noise condition:

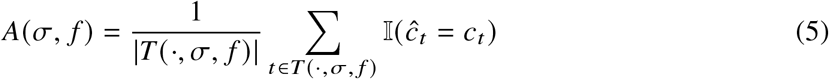

where 𝕝 (·) is the indicator function. This produced an accuracy matrix *A* ∈ [0, 1] ^| Σ|×|*F*|^.

##### 4. Behavioral accuracy matrix

Analogous to the neural decoding matrix, we computed a behavioral accuracy matrix using the psychophysical data from Subramanian et al. (2023) (see github.com/ajaysub110/critical-band-masking). The behavioral accuracy *A*_*behav*_ (*σ, f*) was calculated simply as the proportion of trials for condition (*σ, f*) in which the subject correctly identified the object.

#### Channel fitting

We characterized the spectral tuning of visual cortex by fitting mechanistic models to the summary matrices described above. We utilized distinct modeling approaches for the noise-response and object-decoding data to account for their different signal structures.

##### 1. Modeling the noise-response channel

To quantify the response band, we fit a separable model to the noise-response matrix *R*(*σ, f*), characterizing the BOLD amplitude as a joint function of noise contrast *σ* and spatial frequency *f*. The model assumes that the response is the product of a Naka-Rushton contrast-response function and a Gaussian spatial-frequency tuning curve, plus a baseline:

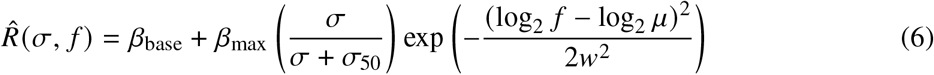

where *μ* and *w* are the center frequency and width of the tuning, *σ*_50_ is the semi-saturation contrast, and *β*_base_ and *β*_max_ scale the response magnitude. We estimated the parameters for each subject and ROI by minimizing the sum of squared errors between the model predictions and the observed BOLD responses. The bandwidth of the noise-response channel was defined as the full width at half maximum (FWHM) of the Gaussian tuning component.

##### 2. Modeling the object-decoding channel

To quantify the recognition band, we fit a probabilistic model to the trial-wise decoding accuracy. We hypothesized that the impairment of object decoding is driven by the strength of the neural response to the noise. Accordingly, the input stage of this model mirrors the structure of the noise-response model described above.

First, the channel’s sensitivity to noise spatial frequency, *S*(*f*), is modeled using the same Gaussian tuning function:

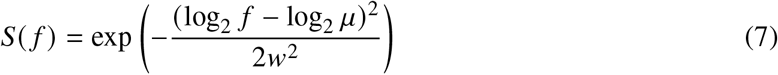

The effective noise drive is the product of this sensitivity and the input noise contrast: *D* (*σ,f*) = *σ* · *S*(*f*). To capture the same gain-control mechanisms observed in the BOLD response, this drive is transformed by a divisive normalization nonlinearity (Heeger, 1993):

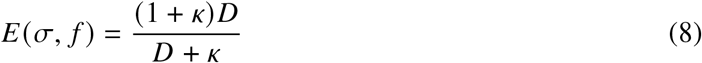

where *E* represents the normalized neural response to the noise and *k* controls the normalization strength.

The key distinction between the two models lies in the final stage. While the noise-response model outputs the response amplitude directly, the decoding model assumes this noise activity interferes with the decision process. We therefore map the normalized noise response *E* to the probability of correct classification using a logistic function:

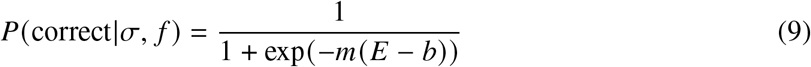

where *m* and *b* determine the slope and threshold of the readout constraint. We fit the model parameters by minimizing the negative log-likelihood of the observed single-trial classification outcomes (correct vs. incorrect), assuming a binomial distribution.

##### 3. Measuring recognition bandwidth

Unlike the response bandwidth, the recognition bandwidth was defined based on the noise power required to impair performance, providing a metric comparable to psychophysical critical-band masking. Using the fitted decoding model, we calculated a *noise power threshold* curve Θ (*f*)—defined as the noise variance (*σ*^2^) required at frequency *f* to reduce decoding accuracy to 50% of the noise-free baseline. We defined the recognition bandwidth as the full width of this threshold curve at twice the minimum power (FW2M). This metric corresponds to the spectral bandwidth at half-power sensitivity (3 dB). The same procedure was applied to the behavioral accuracy matrices to derive the psychophysical recognition bandwidth.

## Footnotes

1 The behavioral bandwidth of 1.5 octaves reported here, refit from Subramanian et al. (2023), differs slightly from the 1-octave estimate in Subramanian et al. (2023) because the channel-fitting procedures differ. Subramanian et al. (2023) fits psychometric functions separately for each noise band to obtain thresholds, and then fit a Gaussian to those thresholds. Here, we use a joint two-dimensional fit over spatial frequency and noise strength, which improves robustness by reducing the number of fitted parameters.

## References

Aghajari, S., Vinke, L. N., and Ling, S. (2020). Population spatial frequency tuning in human early visual cortex. Journal of neurophysiology, 123(2):773–785.

Allen, E. J., St-Yves, G., Wu, Y., Breedlove, J. L., Prince, J. S., Dowdle, L. T., Nau, M., Caron, B., Pestilli, F., Charest, I., et al. (2022). A massive 7t fmri dataset to bridge cognitive neuroscience and artificial intelligence. Nature neuroscience, 25(1):116–126.

Benson, N. C., Jamison, K. W., Arcaro, M. J., Vu, A. T., Glasser, M. F., Coalson, T. S., Van Essen, D. C., Yacoub, E., Ugurbil, K., Winawer, J., et al. (2018). The human connectome project 7 tesla retinotopy dataset: Description and population receptive field analysis. Journal of vision, 18(13):23–23.

Benson, N. C. and Winawer, J. (2018). Bayesian analysis of retinotopic maps. elife, 7:e40224.

Benson, N. C., Yoon, J. M., Forenzo, D., Engel, S. A., Kay, K. N., and Winawer, J. (2022). Variability of the surface area of the v1, v2, and v3 maps in a large sample of human observers. Journal of Neuroscience, 42(46):8629–8646.

Burt, P. J. and Adelson, E. H. (1987). The laplacian pyramid as a compact image code. In Readings in computer vision, pages 671–679. Elsevier.

Campbell, F. W. and Robson, J. G. (1968). Application of fourier analysis to the visibility of gratings. The Journal of physiology, 197(3):551.

Chen, T., Kornblith, S., Norouzi, M., and Hinton, G. (2020). A simple framework for contrastive learning of visual representations. In International conference on machine learning, pages 1597–1607. PmLR.

Cox, R. W. and Hyde, J. S. (1997). Software tools for analysis and visualization of fmri data. NMR in Biomedicine: An International Journal Devoted to the Development and Application of Magnetic Resonance In Vivo, 10(4-5):171–178.

Dale, A. M., Fischl, B., and Sereno, M. I. (1999). Cortical surface-based analysis: I. segmentation and surface reconstruction. Neuroimage, 9(2):179–194.

Deng, J., Dong, W., Socher, R., Li, L.-J., Li, K., and Fei-Fei, L. (2009). Imagenet: A large-scale hierarchical image database. In 2009 IEEE conference on computer vision and pattern recognition, pages 248–255. Ieee.

Dumoulin, S. O. and Wandell, B. A. (2008). Population receptive field estimates in human visual cortex. Neuroimage, 39(2):647–660.

Epstein, R. and Kanwisher, N. (1998). A cortical representation of the local visual environment. Nature, 392(6676):598–601.

Esteban, O., Markiewicz, C. J., Blair, R. W., Moodie, C. A., Isik, A. I., Erramuzpe, A., Kent, J. D., Goncalves, M., DuPre, E., Snyder, M., et al. (2019). fmriprep: a robust preprocessing pipeline for functional mri. Nature methods, 16(1):111–116.

Feinberg, D. A., Moeller, S., Smith, S. M., Auerbach, E., Ramanna, S., Glasser, M. F., Miller, K. L., Ugurbil, K., and Yacoub, E. (2010). Multiplexed echo planar imaging for sub-second whole brain fmri and fast diffusion imaging. PloS one, 5(12):e15710.

Fischl, B. (2012). Freesurfer. Neuroimage, 62(2):774–781.

Fletcher, H. (1940). Auditory patterns. Reviews of modern physics, 12(1):47.

Freeman, J., Ziemba, C. M., Heeger, D. J., Simoncelli, E. P., and Movshon, J. A. (2013). A functional and perceptual signature of the second visual area in primates. Nature neuroscience, 16(7):974–981.

Geirhos, R., Rubisch, P., Michaelis, C., Bethge, M., Wichmann, F. A., and Brendel, W. (2018a). Imagenet-trained cnns are biased towards texture; increasing shape bias improves accuracy and robustness. In International conference on learning representations.

Geirhos, R., Temme, C. R., Rauber, J., Schütt, H. H., Bethge, M., and Wichmann, F. A. (2018b). Generalisation in humans and deep neural networks. Advances in neural information processing systems, 31.

Geisler, W. S. (2011). Contributions of ideal observer theory to vision research. Vision research, 51(7):771–781.

Gorgolewski, K. J., Auer, T., Calhoun, V. D., Craddock, R. C., Das, S., Duff, E. P., Flandin, G., Ghosh, S. S., Glatard, T., Halchenko, Y. O., et al. (2016). The brain imaging data structure, a format for organizing and describing outputs of neuroimaging experiments. Scientific data, 3(1):1–9.

Greve, D. N. and Fischl, B. (2009). Accurate and robust brain image alignment using boundary-based registration. Neuroimage, 48(1):63–72.

Grill-Spector, K. and Weiner, K. S. (2014). The functional architecture of the ventral temporal cortex and its role in categorization. Nature Reviews Neuroscience, 15(8):536–548.

Ha, J., Broderick, W. F., Kay, K., and Winawer, J. (2025). Spatial frequency maps in human visual cortex: A replication and extension. bioRxiv.

Harel, A., Kravitz, D. J., and Baker, C. I. (2014). Task context impacts visual object processing differentially across the cortex. Proceedings of the National Academy of Sciences, 111(10):E962–E971.

Heeger, D. J. (1993). Modeling simple-cell direction selectivity with normalized, half-squared, linear operators. Journal of neurophysiology, 70(5):1885–1898.

Hung, C. P., Kreiman, G., Poggio, T., and DiCarlo, J. J. (2005). Fast readout of object identity from macaque inferior temporal cortex. Science, 310(5749):863–866.

Kay, K. N., Weiner, K. S., and Grill-Spector, K. (2015). Attention reduces spatial uncertainty in human ventral temporal cortex. Current Biology, 25(5):595–600.

Kay, K. N. and Yeatman, J. D. (2017). Bottom-up and top-down computations in word-and face-selective cortex. elife, 6:e22341.

Kleiner, M., Brainard, D., and Pelli, D. (2007). What’s new in psychtoolbox-3?

Kriegeskorte, N., Mur, M., Ruff, D. A., Kiani, R., Bodurka, J., Esteky, H., Tanaka, K., and Bandettini, P. A. (2008). Matching categorical object representations in inferior temporal cortex of man and monkey. Neuron, 60(6):1126–1141.

Kurzawski, J. W., Qiu, B. S., Majaj, N. J., Benson, N. C., Pelli, D. G., and Winawer, J. (2025). Human v4 size predicts crowding distance. Nature communications, 16(1):3876.

Lieber, J. D., Oleskiw, T. D., Simoncelli, E. P., and Movshon, J. A. (2024). Responses of neurons in macaque v4 to object and texture images. BioRxiv, pages 2024–02.

Madry, A., Makelov, A., Schmidt, L., Tsipras, D., and Vladu, A. (2017). Towards deep learning models resistant to adversarial attacks. arXiv preprint arXiv:1706.06083.

Majaj, N. J., Pelli, D. G., Kurshan, P., and Palomares, M. (2002). The role of spatial frequency channels in letter identification. Vision research, 42(9):1165–1184.

Maunsell, J. H. and Treue, S. (2006). Feature-based attention in visual cortex. Trends in neurosciences, 29(6):317–322.

Moeller, S., Yacoub, E., Olman, C. A., Auerbach, E., Strupp, J., Harel, N., and Uğurbil, K. (2010). Multiband multislice ge-epi at 7 tesla, with 16-fold acceleration using partial parallel imaging with application to high spatial and temporal whole-brain fmri. Magnetic resonance in medicine, 63(5):1144–1153.

Neri, P. (2015). The elementary operations of human vision are not reducible to template-matching. PLoS computational biology, 11(11):e1004499.

Peelen, M. V. and Downing, P. E. (2005). Selectivity for the human body in the fusiform gyrus. Journal of neurophysiology, 93(1):603–608.

Pratte, M. S., Ling, S., Swisher, J. D., and Tong, F. (2013). How attention extracts objects from noise. Journal of neurophysiology, 110(6):1346–1356.

Prince, J. S., Charest, I., Kurzawski, J. W., Pyles, J. A., Tarr, M. J., and Kay, K. N. (2022). Improving the accuracy of single-trial fmri response estimates using glmsingle. Elife, 11:e77599.

Radford, A., Kim, J. W., Hallacy, C., Ramesh, A., Goh, G., Agarwal, S., Sastry, G., Askell, A., Mishkin, P., Clark, J., et al. (2021). Learning transferable visual models from natural language supervision. In International conference on machine learning, pages 8748–8763. PmLR.

Rosenke, M., Van den Hurk, J., Margalit, E., Op de Beeck, H., Grill-Spector, K., and Weiner, K. (2020). Extensive individual differences of category information in ventral temporal cortex in the congenitally blind. bioRxiv, pages 2020–06.

Rust, N. C. and DiCarlo, J. J. (2010). Selectivity and tolerance (“invarianceff) both increase as visual information propagates from cortical area v4 to it. Journal of Neuroscience, 30(39):12978–12995.

Schwarzlose, R. F., Baker, C. I., and Kanwisher, N. (2005). Separate face and body selectivity on the fusiform gyrus. Journal of Neuroscience, 25(47):11055–11059.

Solomon, J. A. and Pelli, D. G. (1994). The visual filter mediating letter identification. Nature, 369(6479):395–397.

Stigliani, A., Weiner, K. S., and Grill-Spector, K. (2015). Temporal processing capacity in high-level visual cortex is domain specific. Journal of Neuroscience, 35(36):12412–12424.

Subramanian, A., Sizikova, E., Majaj, N., and Pelli, D. (2023). Spatial-frequency channels, shape bias, and adversarial robustness. Advances in neural information processing systems, 36:4137–4149.

Szegedy, C., Zaremba, W., Sutskever, I., Bruna, J., Erhan, D., Goodfellow, I., and Fergus, R. (2013). Intriguing properties of neural networks. arXiv preprint arXiv:1312.6199.

Weiner, K. S., Barnett, M. A., Witthoft, N., Golarai, G., Stigliani, A., Kay, K. N., Gomez, J., Natu, V. S., Amunts, K., Zilles, K., et al. (2018). Defining the most probable location of the parahippocampal place area using cortex-based alignment and cross-validation. Neuroimage, 170:373–384.

Weiner, K. S., Golarai, G., Caspers, J., Chuapoco, M. R., Mohlberg, H., Zilles, K., Amunts, K., and Grill-Spector, K. (2014). The mid-fusiform sulcus: a landmark identifying both cytoarchitectonic and functional divisions of human ventral temporal cortex. Neuroimage, 84:453–465.

Winawer, J. and Witthoft, N. (2015). Human v4 and ventral occipital retinotopic maps. Visual neuroscience, 32:E020.

Xu, J., Moeller, S., Auerbach, E. J., Strupp, J., Smith, S. M., Feinberg, D. A., Yacoub, E., and Uğurbil, K. (2013). Evaluation of slice accelerations using multiband echo planar imaging at 3 t. Neuroimage, 83:991–1001.

Ziemba, C. M., Freeman, J., Movshon, J. A., and Simoncelli, E. P. (2016). Selectivity and tolerance for visual texture in macaque v2. Proceedings of the National Academy of Sciences, 113(22):E3140–E3149.

