## Supplementary material for "An increasingly efficient narrowband object-recognition channel along the ventral stream"

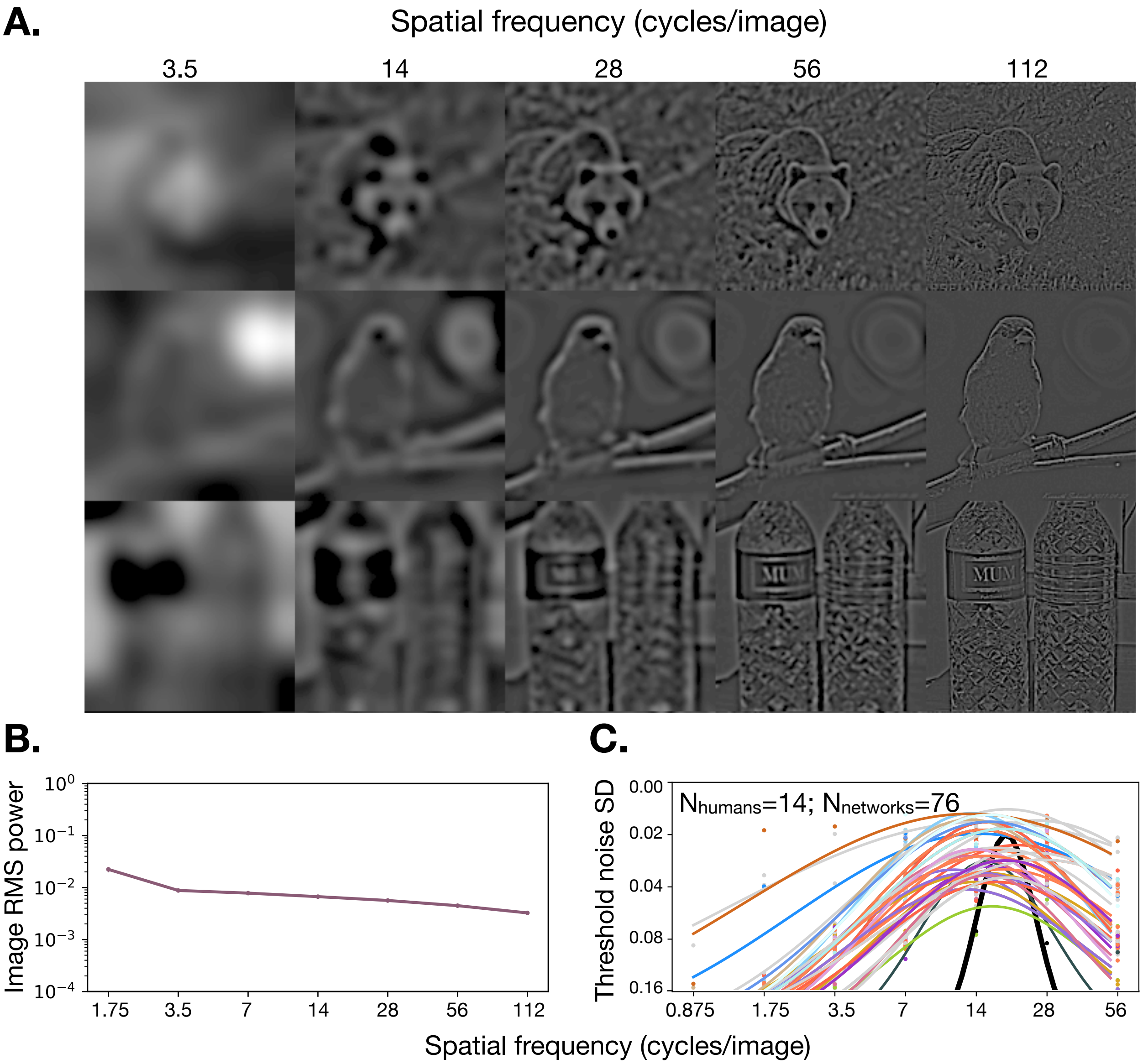


**Figure S1: Three ways of showing that the 1.5 octave object recognition channel is a property of the human visual system, not the stimulus. A.** Visual demonstration of information availability. Significant object information remains visible in off-channel low spatial frequencies (1.75–7 cycles/image), even though human recognition critically relies on the higher channel frequencies (14–112 cycles/image). **B.** Image statistics versus channel tuning. The average RMS power spectrum of the natural image set (purple line) contains significant energy across the entire spectrum, whereas the psychophysical recognition channel (black curve) is restricted to a narrow fraction of these frequencies. **C.** Human versus machine vision. Unlike humans, deep neural networks utilize a broad range of spatial frequencies—including lower frequencies—to categorize images (data from Subramanian et al. (2023)). This divergence suggests the narrow human channel is a specific biological constraint rather than a feature of the image statistics. Note that we use slightly different spatial-frequency ranges for psychophysics (0.875 to 56 cpi) and fMRI (1.75 to 112 cpi).

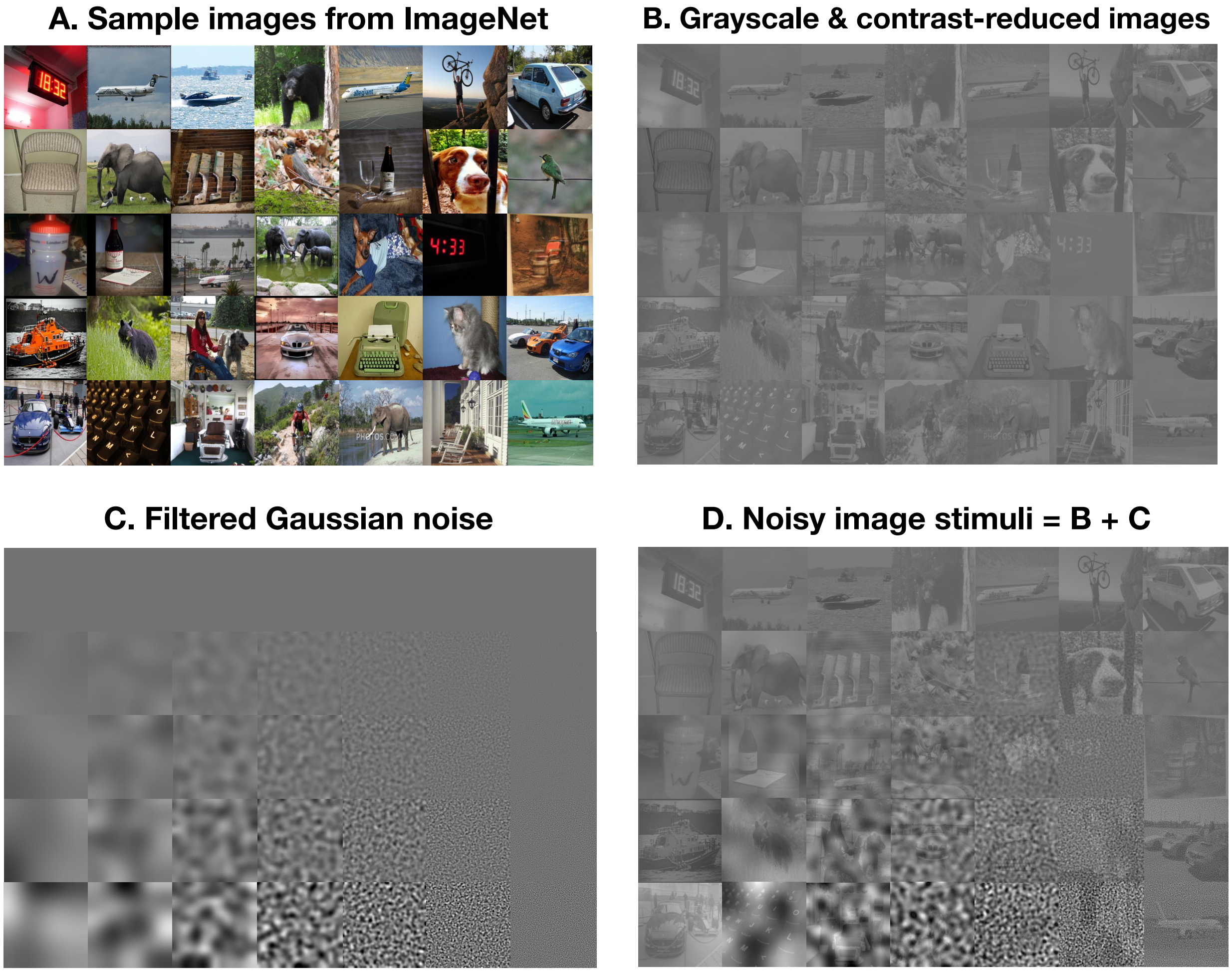


**Figure S2: Stimulus generation pipeline for critical-band masking.**
**A.** Sample natural images drawn from the ImageNet dataset. **B.** Images are converted to grayscale and root-mean-square (RMS) contrast-reduced to 20% of their original intensity to prevent pixel clipping during noise addition. **C.** Samples of filtered Gaussian noise images at each noise contrast (top to bottom) and spatial-frequency band (left to right). **D.** Representative noisy image stimuli used in the fMRI experiment, generated by the additive superposition of the contrast-reduced images (B) and the filtered noise (C).

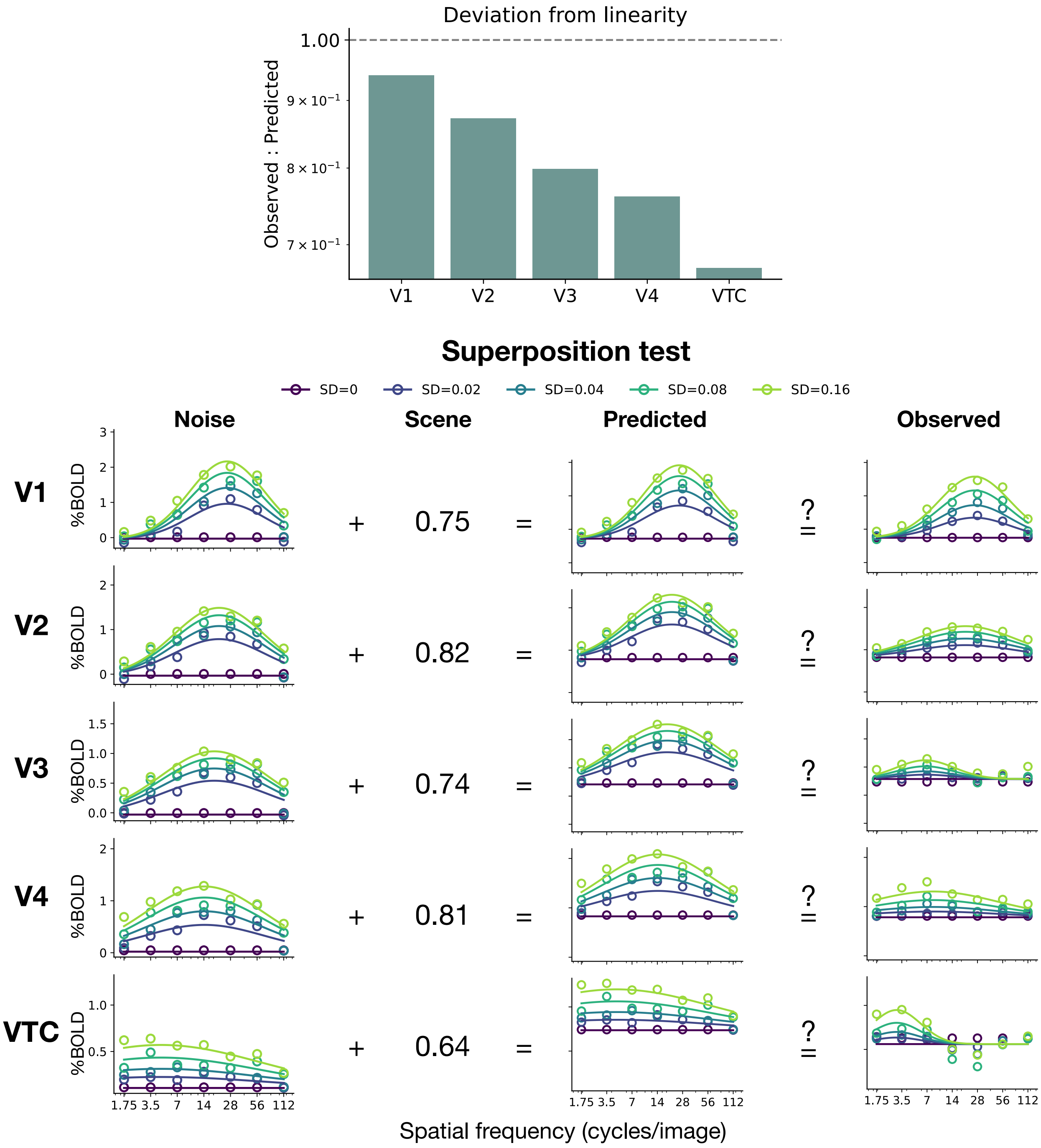


**Figure S3: Superposition test reveals increasing nonlinearity along the ventral stream. Top:** Summary of deviation from linearity. The bar plot displays the ratio of the response to noisy scenes to sum of response to noise and scene. A value of 1.0 (dashed line) indicates perfect linearity. V1 is near-linear, whereas extrastriate areas (V2–VTC) are progressively less linear. **Bottom:** Superposition test by visual area. Rows correspond to visual areas V1 through VTC.

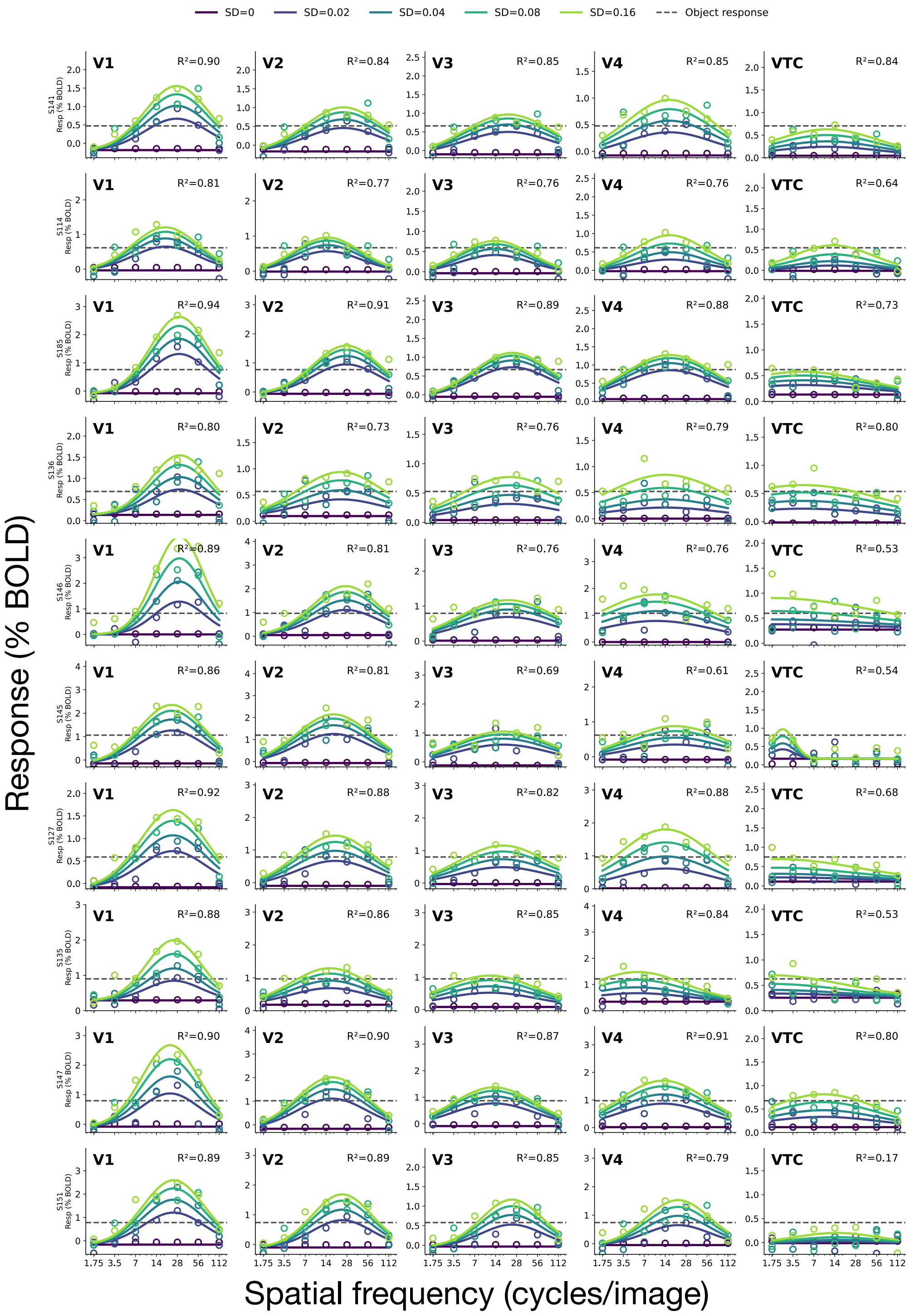

**Figure S4: Measured and predicted noise response for all subjects in all considered visual ROIs.** Each panel shows the individual noise-response channel fit for one participant across the visual hierarchy. These plots show that the patterns evident in the average are also evident in the individuals.

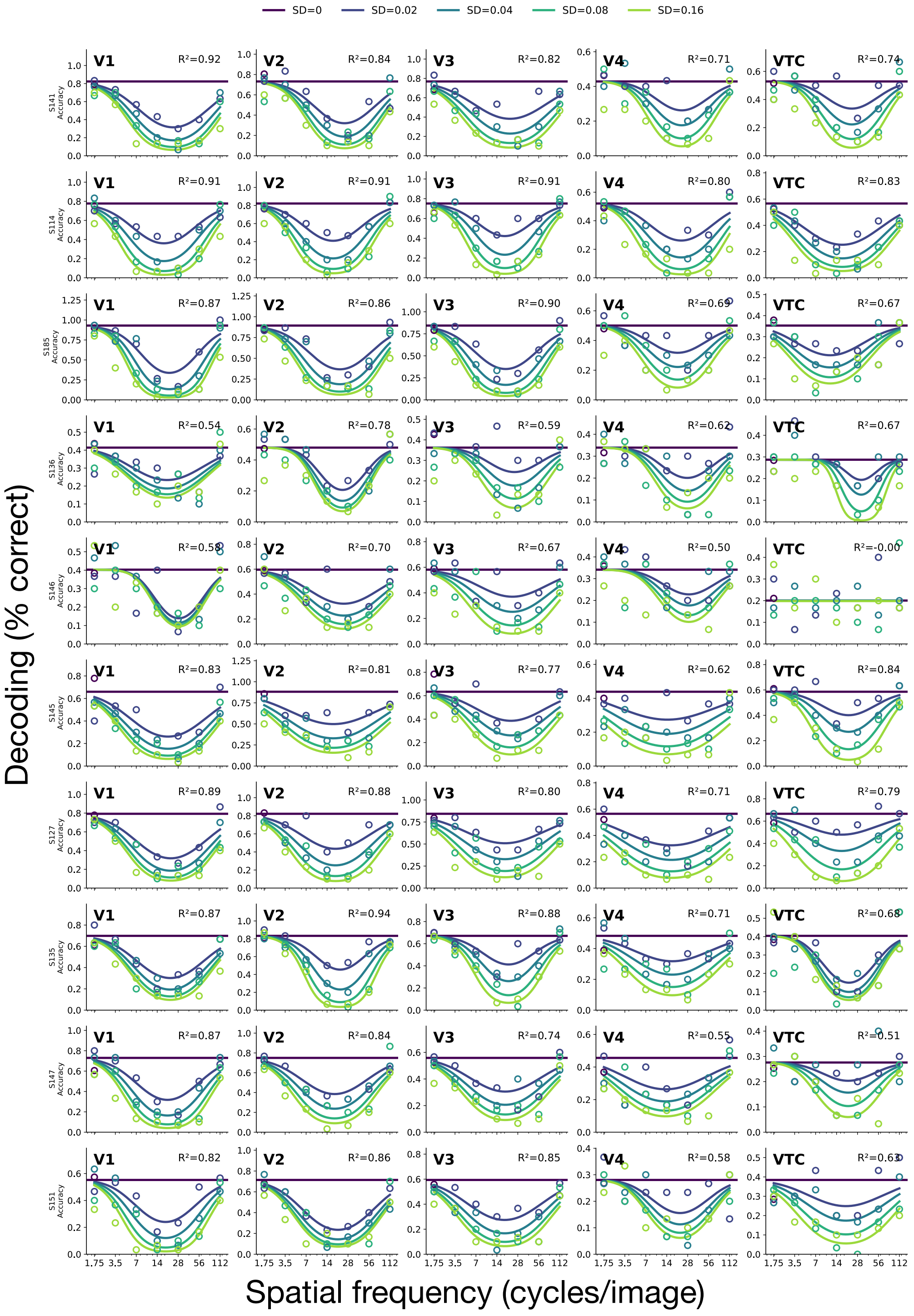


**Figure S5: Measured and predicted object-decoding accuracy for all subjects in all considered visual ROIs.** Individual data points represent raw decoding performance, while solid lines indicate the logistic readout model fits. These plots show that the patterns evident in the average are also evident in the individuals.

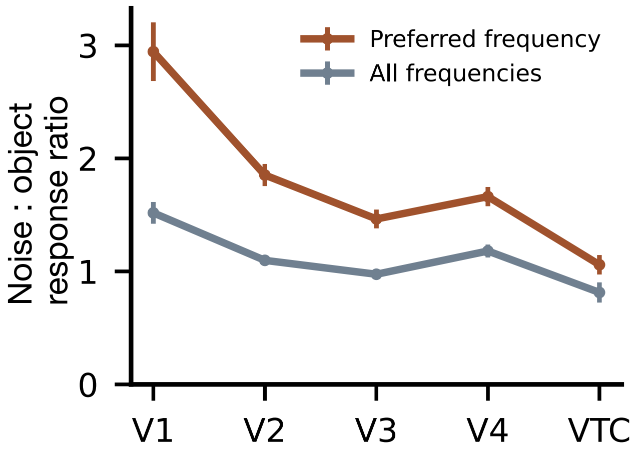


**Figure S6: Ratio between the noise and scene response along the ventral pathway.** Noise response is taken at the highest noise level ($\sigma = 0.16$). Brown line shows ratio when noise response is taken at the preferred frequency. Gray line shows ratio when noise response is averaged across all frequency bands.

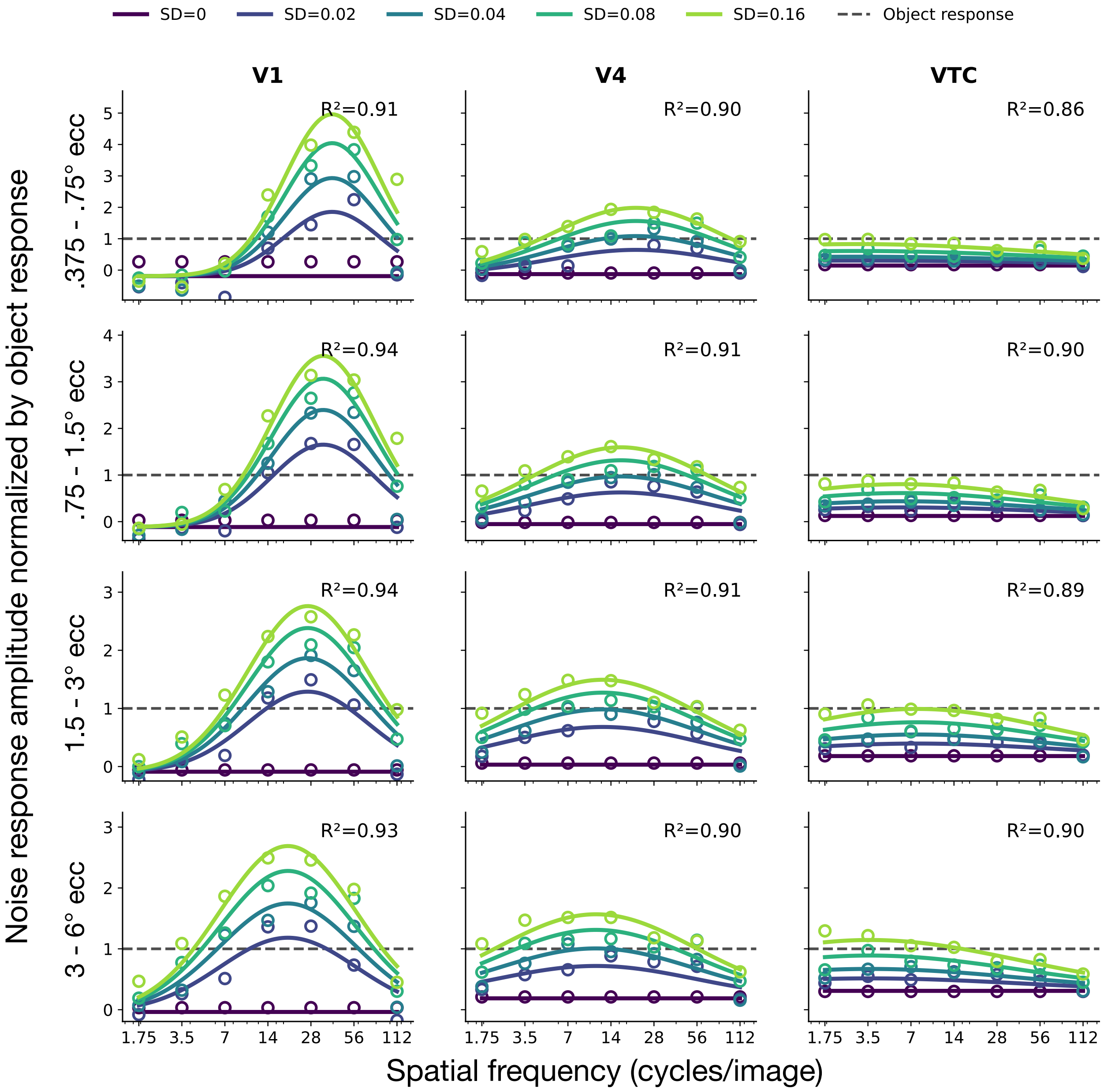


**Figure S7: The noise-response band by eccentricity**. In each area and each eccentricity band, responses are normalized to the scene response (dashed lines). These are the same data as in Figure 2A but separated by eccentricity band. As expected, for V1 (left column), the preferred spatial frequency is highest near the fovea (top row) and decreases with eccentricity (successive rows). The same pattern holds for V4 (middle column). The VTC responses (right column) are low pass across all eccentricities. Overall the figure shows that the patterns in the main text are conserved across eccentricity: a shift from band pass to low pass from V1 to VTC, and a weaker response to noise from V1 to VTC, relative to the scene response.

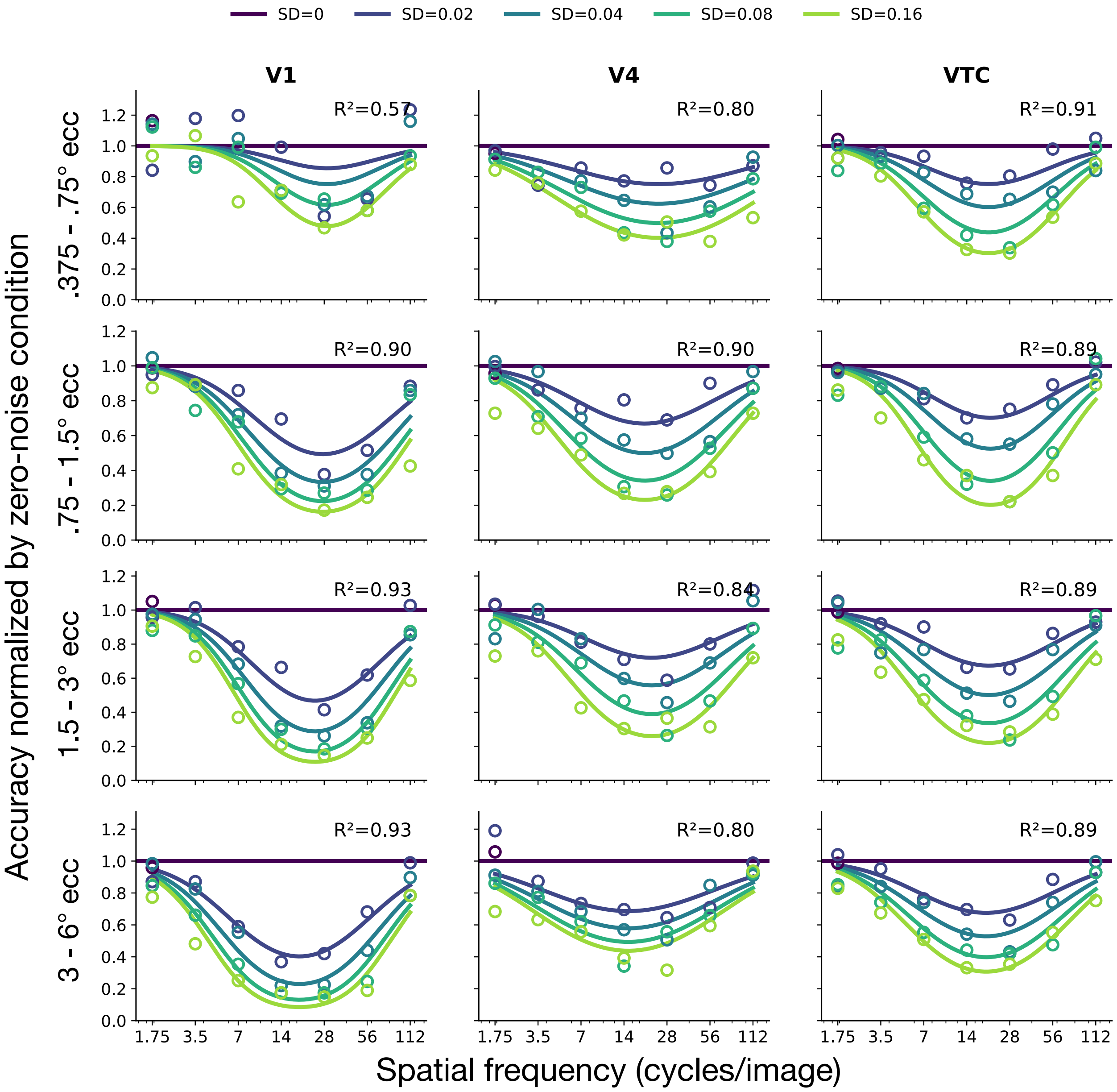


**Figure S8: The recognition band by eccentricity**. In each area and each eccentricity band, decoding accuracy is normalized to zero-noise condition (dark purple lines). These are the same data as in Figure 2B but separated by eccentricity band. In each area and each eccentricity band, the recognition band is narrow. Unlike the response band (Figure S7), the center frequency doesn't shift systematically with eccentricity.
